# Substrate specificity of SARS-CoV-2 nsp10-nsp16 methyltransferase

**DOI:** 10.1101/2020.07.30.228478

**Authors:** Roberto Benoni, Petra Krafcikova, Marek R. Baranowski, Joanna Kowalska, Evzen Boura, Hana Cahová

**Affiliations:** Institute of Organic Chemistry and Biochemistry of the Czech Academy of Sciences, 16610 Prague, Czech Republic; Division of Biophysics, Institute of Experimental Physics, Faculty of Physics, University of Warsaw, Ludwika Pasteura 5, 02-093 Warsaw

**Keywords:** virus, SARS-CoV-2, methylation, inhibitor

## Abstract

The ongoing COVID-19 pandemic exemplifies the general need to better understand viral infections. The positive single strand RNA genome of its causative agent, the SARS coronavirus 2 (SARS-CoV-2) encodes all viral enzymes. In this work, we focus on one particular methyltransferase (MTase), nsp16, which in complex with nsp10 is capable of methylating the first nucleotide of a capped RNA strand at the 2′-O position. This process is part of a viral capping system and is crucial for viral evasion of the innate immune reaction. In light of recently discovered non-canonical RNA caps, we tested various dinucleoside polyphosphate-capped RNAs as substrates for nsp10-nsp16 MTase. We developed an LC-MS-based method and discovered five types of capped RNA (m^7^Gp_3_A(G)-, Gp_3_A(G)- and Gp_4_A-RNA) that are substrates of the nsp10-nsp16 MTase. Our technique is an alternative to the classical isotope labelling approach for measurement of 2′-O-MTase activity. Further, we determined the IC_50_ value of sinefungin (286 ± 66 nM) to illustrate the value of our approach for inhibitor screening. In the future, this approach can be used for screening inhibitors of any type of 2′-O-MTase.

## Introduction

The severe acute respiratory syndrome coronavirus 2 (SARS-CoV-2) is the causative agent of the current COVID-19 pandemic [1] that has already infected more than ten million human beings and claimed over 600 thousand lives according to the World Health Organization (WHO, www.who.int). It belongs to the *Coronaviridae* family that has already produced at least two other deadly human viruses during the last two decades. The severe acute respiratory syndrome (SARS) virus was identified as the virus causing atypical pneumonia in the Guangdong Province of China in 2002 [2] and the Middle East Respiratory Syndrome (MERS) virus was responsible for the outbreak of a respiratory disease in 2012 in the Arabian Peninsula region [3].

Coronaviruses are now recognized as a major threat to global human health [4]. Their genome is a single-stranded positive sense RNA that encodes four structural and sixteen non-structural (nsp1-16) proteins [5]. It is the non-structural proteins that perform all enzymatic activity essential for the viral lifecycle that are not available in the host cell. Those are the RNA-dependent RNA-polymerase (RdRp); the two proteases, papain-like protease (PL^pro^) and 3C-like main proteases (3CL^pro^); the nsp13 helicase and two methyltransferases [5]. Each of these enzymes is a potential target for antivirals [6] and SARS-CoV-2 enzymes are therefore intensively studied. The prime target is the RdRp, a heterotrimeric protein complex composed of nsp7, nsp8, and nsp12. The only small molecule currently approved for experimental treatment by the FDA, remdesivir, inhibits the RdRp [7]. The RdRp was well structurally characterized including its interaction with RNA and with remdesivir [8-11]. Also the structure and first inhibitors of the main protease 3CL^pro^ were recently described [12] and the first structures of MTases were solved [13-15].

Innate immunity is a crucial part of the human immune system and viruses have evolved abilities to evade it [16]. The 5′-end of the nascent RNA is a part of the pattern recognized by the RIG-I (retinoic acid-inducible gene I) pattern recognition receptor. It recognizes short viral dsRNA with a 5′-triphosphate [17] or 5′-diphosphate [18] which leads to interferon (IFN) expression. Subsequently IFN-induced proteins with tetratricopeptide repeats 1 and 5 (IFIT 1 and IFIT5) sequester uncapped (5′-triphosphorylated) and 5′-capped RNAs lacking 2′-O-methylation at the first transcribed nucleotide (RNA carrying cap-0) which prevents binding to the eukaryotic translation initiation factor 4E (EIF4E) and inhibits its translation [19]. Coronaviruses have two RNA MTases, nsp14 and nsp16, that ensure the creation of the RNA cap (Figure 1). Nsp14 is an N7-MTase that methylates the first GTP nucleobase and, subsequently, nsp16, a 2′-O-MTase methylates the following nucleotide. Interestingly the SARS-CoV nsp16 is only active when it is in complex with nsp10 that acts as its activation factor [20].

**Figure 1:**
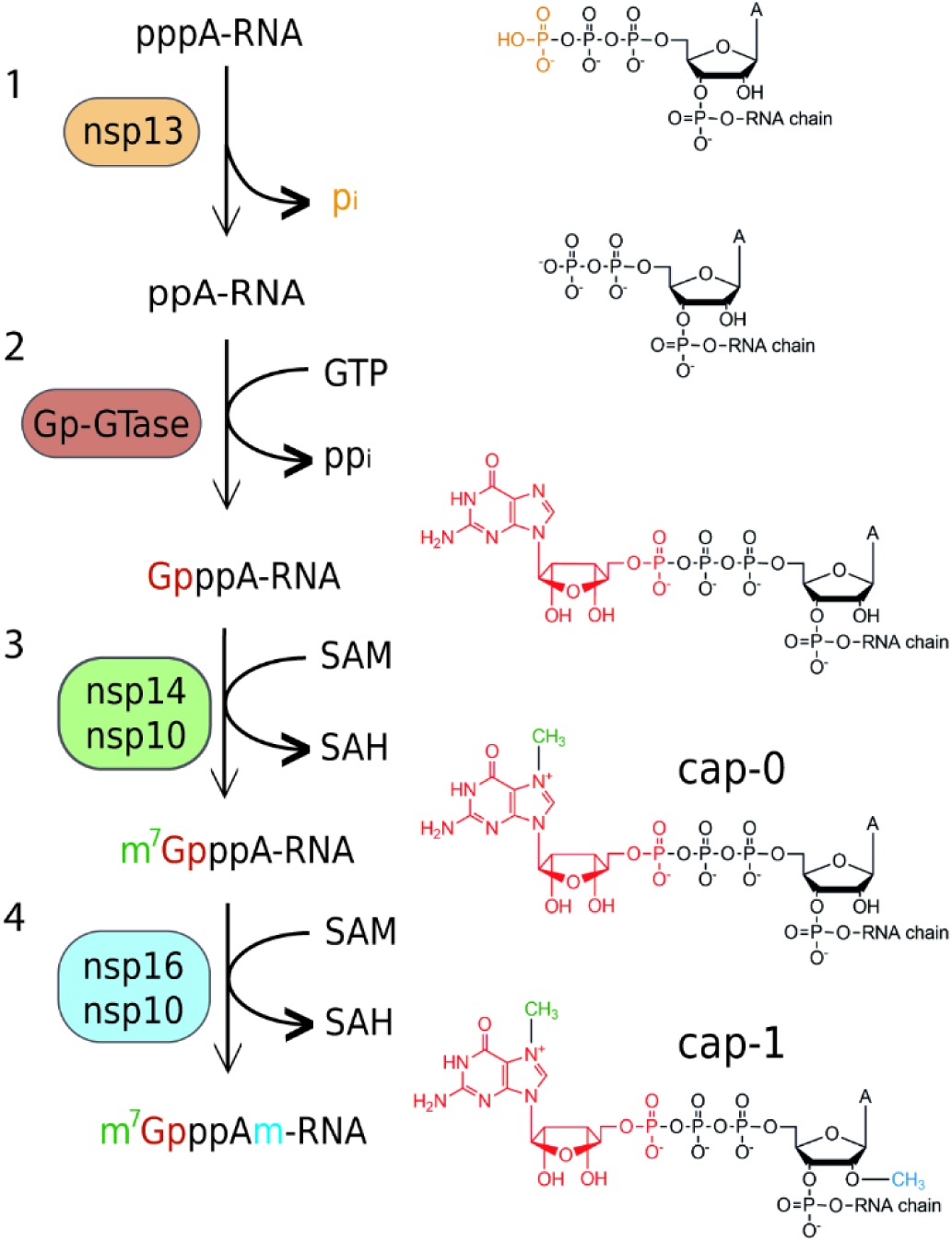
Overview of cap 1 structure formation in SARS-CoV-2: *i*). The hydrolysis of the 5′γ-phosphate of the nascent RNA (pppA-RNA) by an RNA 5′-triphosphatase (nsp13 helicase). *ii*) An unknown guanylyltransferase (GTase) in a two-step reaction transfers GMP to form the cap core structure (GpppA). *iii*) nsp14 methyltransferase with a co-factor nsp10 methylates guanosine at the N7 position and forms the cap-0 structure (m7GpppA). *iv*) Nsp16 in complex with nsp10 methylates ribose at the 2′O position of the first transcribed nucleotide to form the cap-1 structure (m7GpppAm).

The chemical variations in RNA caps and their physiological implications are not fully understood. Recently, it has been shown that beside the common canonical m^7^Gp_3_N cap, RNA can be capped by cofactors such as nicotinamide adenine dinucleotide [21, 22] or coenzyme A [23, 24]. While the regulatory role of the NAD-cap in bacteria has been partially elucidated [25], its function in mammalian cells has not been fully understood yet [26], albeit it was suggested that it promotes RNA decay [22]. The role of the CoA-cap is unknown. Recently, we reported the discovery of an entirely new class of 5′ RNA caps in bacteria [27]. These caps have the structure of dinucleoside polyphosphates (Np_*n*_Ns) and are incorporated into RNA co-transcriptionally by the RNA polymerase [28]. Dinucleoside polyphosphates have been known for more than 50 years and have been detected in all kingdoms of life, including human cells [29]. They are often called alarmones, as their intracellular concentration increases under stress condition [30]. As Np_*n*_Ns are also present also in eukaryotic cells, we hypothesize that they might be incorporated into RNA as non-canonical initiating nucleotides where they can represent an additional layer of information. Moreover, NAD or flavin adenine dinucleotide (FAD) capped RNA was detected in viral particles of Dengue 2 virus [31], suggesting that non-canonical RNA caps might play a role in the viral life cycle.

So far, RNA capped with non-canonical initiating nucleotides such as NAD, CoA or Np_*n*_Ns have not been studied as substrates for any viral encoded enzyme.

Here, we aimed to characterize the SARS-CoV-2 nsp10-nsp16 2′-O-MTase. We prepared a recombinant nsp10-nsp16 complex and analysed its substrate specificity using LC-MS. First, we tested whether nsp10-nsp16 is capable of methylation of free caps or short hexamer RNA capped with canonical and non-canonical RNA nucleotides. As we did not observe any methylation of the free caps and the methylation of the short hexamer RNA was only partial, we used a longer RNA (35mer). Usually, the methylation of RNA at the 2′-O of ribose is studied by radioactive labelling [20]. We developed a new general technique that can be used for the analysis of any cellular or viral RNA MTase. RNA prepared bearing various caps *in vitro* is treated with an MTase and then digested by the Nuclease P1 into nucleotides and caps. The efficiency of the reaction is followed by LC-MS analysis of digested RNA before and after methylation reactions. Our analysis showed that nsp10-nsp16 2′-O-MTase can methylate ribose at the 2′ position of RNA capped with m^7^Gp_3_A, Gp_3_A, m^7^Gp_3_G, Gp_3_G and Gp_4_A. We discovered that the m^7^Gp_3_A-RNA was the best substrate for nsp10-nsp16 in accordance with studies on MTases from other coronaviruses [20, 32, 33]. We also show that this method is suitable for characterization of MTases inhibitors. As a model compound, we used the pan-MTase inhibitor sinefungin [34] and we obtained an IC_50_ value of 286 ± 66 nM.

## Results and Discussion

### Methyltransferase complex of nsp10-nsp16 does not methylate free RNA caps

In the light of our recent discovery of a new class RNA caps based on dinucleoside polyphosphates (Np_*n*_Ns) [27], we tested whether nsp16/nsp10 may methylate 2′-O position of ribose from various Np_*n*_Ns. We let m^7^Gp_3_A, Gp_3_A, Ap_3_A, m^7^Gp_3_G, Gp_3_G, Np_4_N (N=A, G) react with nsp10-nsp16 complex in the presence of SAM for 2 h at 30 °C or 37 °C. The reaction mixture was analysed by HPLC. We did not observe any 2′-O-methylated products. This finding was in an agreement with previously observed SARS-CoV nsp10-nsp16 activity [20] (Figure S1).

### Methyltransferase complex of nsp10-nsp16 partially methylates the short m^7^Gp_3_A-RNA

We also tested whether a short RNA (6mer) capped with various dinucleoside polyphosphates can be methylated by this complex. The hexameric RNA was prepared by *in vitro* transcription with T7 RNA polymerase and free caps. After HPLC purification, RNA was treated by nsp10-nsp16 complex with SAM for 2 h at 30 °C. The samples were then digested by the nuclease P1 to release 5’-mononucleotides and intact RNA caps and analysed by HPLC. From all the tested substrates (m^7^Gp_3_A-, Gp_3_A-, NAD-RNA) only m^7^Gp_3_A-RNA was methylated in approximately 20 % yield (Figure S2). This experiment showed that the activity of the complex can be observed once a hexameric RNA is used. Although for the development of an inhibitor screening assay another approach with higher enzymatic activity is desired.

### LC-MS method for the methyltransferase activity of nsp10-nsp16

Since the hexamer RNA was not an ideal substrate for nsp10-nsp16, we prepared a 35mer RNA with m^7^Gp_3_A cap by *in vitro* transcription and treated it with nsp10-nsp16 complex and SAM at 30 °C for 30 min, 1 h and 2 h. After indicated times, the samples were digested by Nuclease P1 and analysed by LC-MS [27]. We followed the disappearance of the unreacted cap (m^7^Gp_3_A) and observed the formation of 2′-O-methylated m^7^Gp_3_A (m^7^Gp_3_A_m_). After 2 h, all m^7^Gp_3_A cap was converted to m^7^Gp_3_A_m_. We choose these conditions for the following screening of other capped-RNAs.

We tested thirteen differently capped RNAs in total (m^7^Gp_3_A, m^6^Ap_3_A, m^7^Gp_3_G, Ap_3-5_N, Gp_3-4_G, NAD, CoA) as a substrate for the SARS-CoV-2 nsp10-nsp16 MTase complex. The RNA was prepared as a 35mer by *in vitro* transcription, treated by the nsp10-nsp16 complex in the presence of SAM at 30°C for 2 h. Afterwards, the samples were digested by nuclease P1 and the disappearance of the unreacted cap and formation of the methylated strand was observed (Figure 2A). The efficiency of the enzyme activity was calculated by disappearance of the unreacted cap (Figure 2B). The values were normalized using guanosine monophosphate (GMP) area under the curve (AUC). Under the conditions optimized for m^7^Gp_3_A-RNA, four other capped RNAs (Gp_3_A-, Gp_3_G-, m^7^Gp_3_G- and Gp_4_A-RNA) were methylated at the 2′-O position of the +1 nucleotide. All of them were methylated approximately from 50 % to 10 % (Figure 2C, Figure S3-7) in comparison with m^7^Gp_3_A-RNA. When Ap3G was incorporated into RNA in the opposite manner [28], i.e. A is flanking, such capped RNA was not accepted as substrate of nsp10-nsp16 MTase at all. Besides Np_*n*_Ns-RNA, which have not been detected in eukaryotic cells so far, we also tested the recently discovered eukaryotic NAD-[22] and CoA-RNA [24] as substrates for the nsp10-nsp16 MTase. Even though the NAD cap has a positive charge similar to that of the canonical m^7^Gp_3_A cap, we did not observe any methylated products. Ap_3-5_A-, m^6^Ap_3_A-, Gp5A-, Gp_4_G-, m^7^Gp_4_G-, and CoA-RNA were not accepted as substrates either. In general, the common pattern shared by all methylated substrates is a polyphosphate bridge with 3 to 4 phosphates and a flanking G (Figure 3). Moreover, methylation at the N7 position of G led to a higher yield of 2′-O methylation of the +1 nucleotide, both m^7^Gp_3_A-RNA and m^7^Gp_3_G-RNA were better substrates for the nsp10-nsp16 MTase than their non-methylated counterparts Gp_3_A-RNA and Gp_3_G-RNA (Figure 2C). This finding is in a good agreement with observations on other coronaviruses, showing that the methylation at the position N7 of the flanking guanosine occurs first and the 2′-O methylation at position +1 follows as the second step.

**Figure 2:**
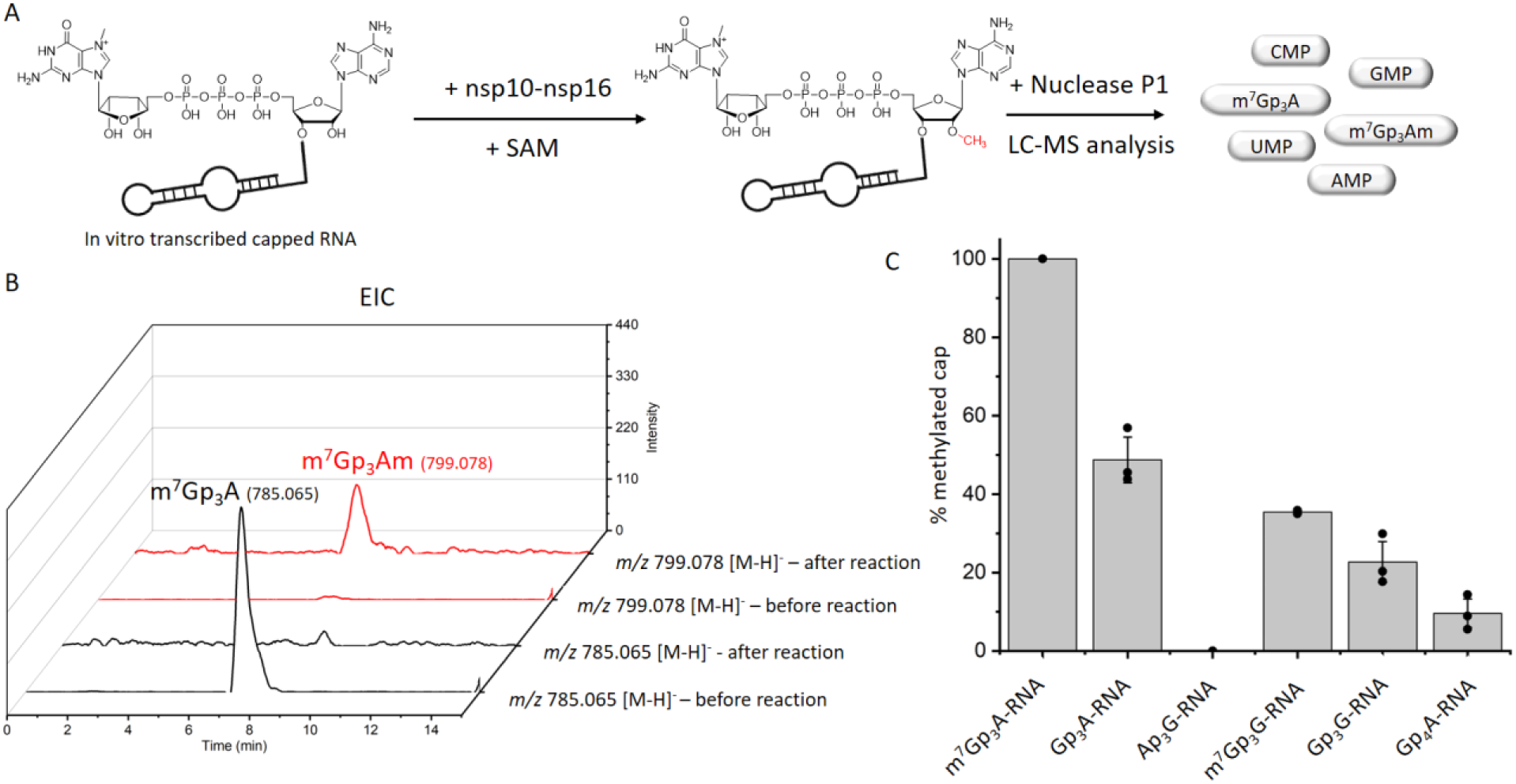
Screening of nsp10-nsp16 activity on non-canonical capped-RNA. A) The scheme of experimental set-up. RNA transcribed in vitro was treated by nsp10-nsp16 and SAM, then treated by nuclease P1 and analysed by LC-MS. B) Extracted Ion Chromatogram (EIC) for m/z 785.065 and m/z 799.078 before and after reaction with nsp10-nsp16. C) The comparison of nsp10-nsp16 methylation efficiency of various capped-RNAs. All measurements were performed in triplicate.

**Figure 3:**
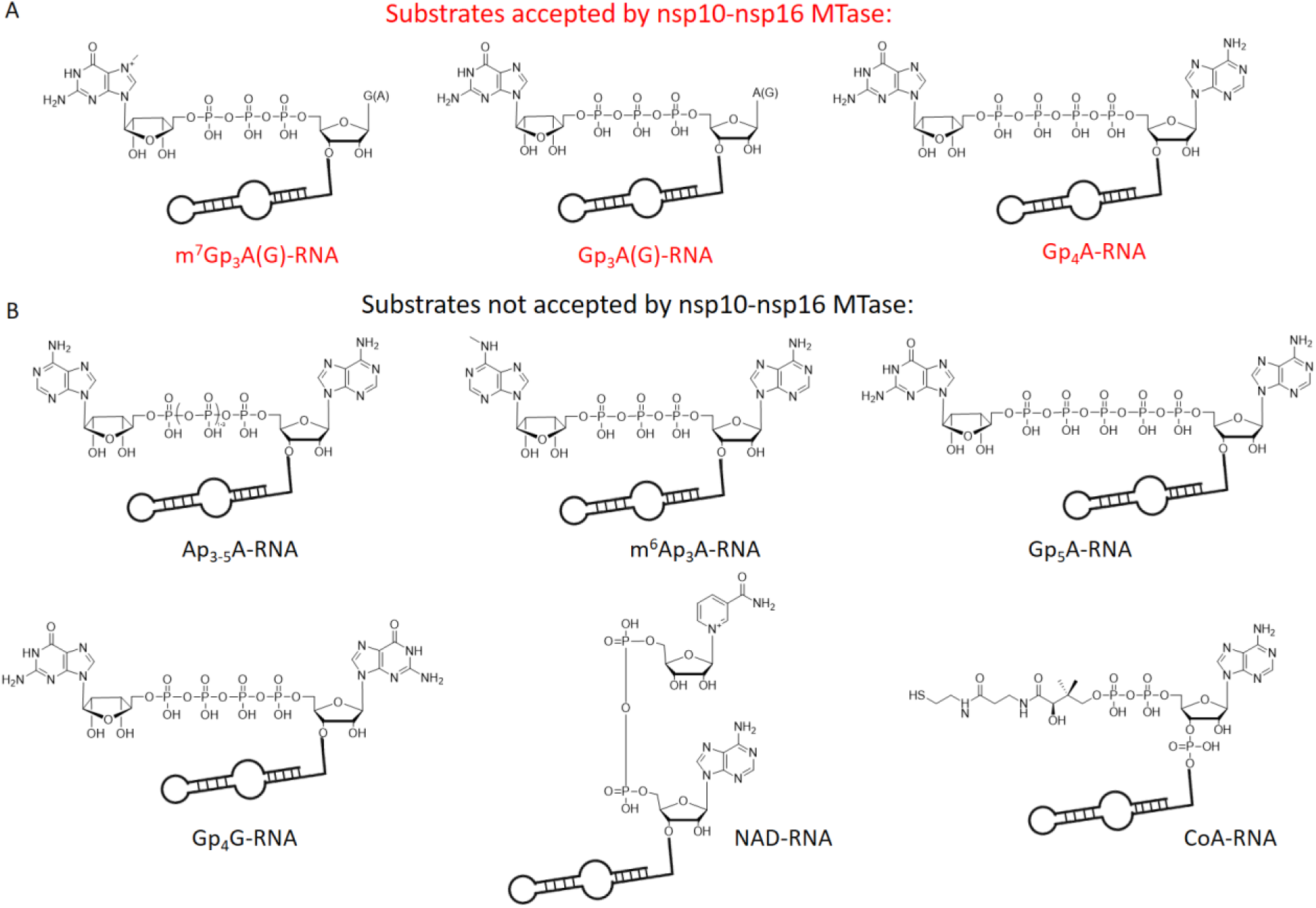
Chemical structures of tested capped-RNAs as substrates of nsp10-nsp16 MTase. A) Substrates accepted by nsp10-nsp16 MTase: m^7^Gp_3_A(G)-, Gp_3_A(G)- and Gp_4_A-RNA. B) Substrates not accepted by nsp10-nsp16 MTase: Ap_3-5_A-, m^6^Ap_3_A-, Gp_5_A-, Gp_4_G-, NAD- and CoA-RNA.

### Non-radioactive LC-MS method for testing of nsp10-nsp16 inhibitors

So far, the methods used for the screening of inhibitors of RNA MTases were based on radioactive labelling. Here, we took an alternative approach and we developed a LC-MS based method for assessing the IC_50_ values of the nsp10-nsp16 MTase inhibitors. Our method is general and can be applied to any RNA MTase and RNA of any sequence. We prepared the m^7^Gp_3_A-RNA substrate *in vitro* and treated it with the nsp10-nsp16 MTase in the presence of SAM and various concentration of the inhibitor. As a model inhibitor, we chose the pan-MTase inhibitor Sinefungin [35]. We optimized the MTase reaction conditions to reach half conversion of the starting capped-RNA. The LC-MS was performed in a positive mode to ensure higher sensitivity of the measurement. Using this method, we were able to determine the IC_50_ value of Sinefungin as 286 ± 66 nM (Figure 4). This value is in a good agreement with the previously published value (736 ± 71 nM) for the SARS-CoV nsp10-nsp16 MTase obtained by a filter-binding assay [20].

**Figure 4:**
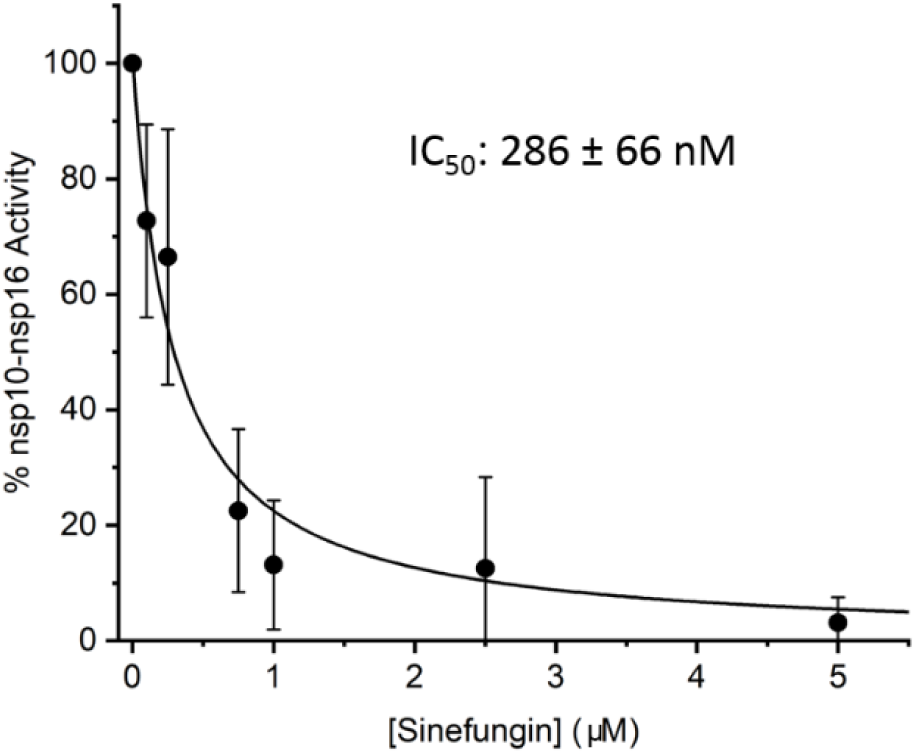
Inhibition curve of Sinefungin. Capped m^7^Gp_3_A-RNA was treated with nsp10-nsp16 and SAM at various concentrations of Sinefungin. After reaction, RNA was cleaved by nuclease P1, analysed and the dimethylated cap (m^7^Gp_3_Am) was quantified by LC-MS. The measurement was performed in triplicate.

## Discussion

Here, we report on the development of an LC-MS-based method for analysis of RNA methylation. Our method is non-radioactive which is the current trend for safety reasons and also advantageous for high throughput screening [36, 37]. We applied our method to the nsp16 MTase from the SARS-CoV-2 to characterize this important drug target. In total, we tested fourteen differently capped RNAs to characterize the substrate specificity of nsp16. As expected, based on the similarity to the SARS-CoV nsp16, the best substrate was m^7^Gp_3_A-RNA [20]. However, we observed that RNAs modified with different caps can also be efficiently methylated: Gp_3_A – 49 %, Gp_3_G – 23 %, m^7^Gp_3_G - 36 % and Gp_4_A - 10 %. This is, surprisingly, not in contradiction to results obtained on coronaviral MTases because previous studies on the SARS-CoV nsp16 used a short (5mer) RNAs that can be methylated only when m^7^Gp_3_A capped [20] which we observed as well when using short 6mer RNA (Figure S1). This has important implication for the viral life cycle. Here we show that RNA that is not yet methylated by the nsp14 N7 MTase can be also a substrate for the 2’-O nsp16 MTase albeit not as good substrate. Nevertheless, this observation challenges the dogma of step-by-step methylation process of coronaviral RNA (Figure 1). Interestingly, the observation of four different caps (Gp_3_A, Gp_3_G, m^7^Gp_3_G and Gp_4_A) also accepted as a substrate for the nsp16 MTase could also play a regulatory role in the stability of viral RNAs. Coronaviruses produce besides the ∼30 kb genomic RNA (serves as mRNA for nsp1-16 proteins) also up to ten subgenomic mRNAs that encode structural and accessory proteins [5]. It would be tempting to speculate that methylation of the subgenomic RNAs could serve a regulatory role and control expression of coronaviral structural and accessory proteins, however, that is unlikely because it was reported that each positive sense SARS-CoV-2 RNA starts with the same 5’ leader sequence [38]. However, various caps can be on an identical sequence. For several polymerases it was shown that if NpnNs are in the proximity of the RNA polymerase, then it accepts them as non-canonical initiating nucleotides [28]. So far, we do not know, if that is also the case for the coronaviral RdRp.

The SARS-CoV-2 nsp16 MTase is an important drug target. Often, drug-like candidate molecules are found using high throughput screening (HTS) [39] and subsequently optimized using medicinal chemistry. Our LC-MS-method could be easily optimized for HTS using a robotic pipeline and small analytical high throughput LC-MS instruments [40, 41] providing a new tool for drug discovery against COVID-19.

Taken together, our LC-MS based approach and an in-depth analysis showed that SARS-CoV-2 nsp16 has a broader substrate specificity than previously believed. Especially the ability of nsp16 to use a non-methylated Gp_3_A has important implications for the viral life cycle because it reveals that nsp16 can, in principle, act before the nsp14 N7 MTase.

## Material and methods

### General

All chemicals were either purchased from Merck or Jena Biosciences and used without further purification. Oligonucleotides were purchased from Generi Biotech. m^7^GpppA was synthesized in house according to Baranowski et al. [42] as detailed in Supplementary Methods.

### Protein expression and purification

The plasmid encoding for nsp10 and nsp16 proteins was described previously as was the purification protocol [13]. Briefly, the expression vector was transformed into *E.coli* BL21 cells and the cells were grown at 37°C in LB media supplemented with 25 μM ZnSO_4_ until the OD_600_ nm reached 0.5. Subsequently, the expression was induced by IPTG (final concentration 300 μM) and the temperature lowered to 18°C overnight. Cells were harvested, resuspended, and lysed by sonication in lysis buffer (50 mM Tris, pH 8, 300 mM NaCl, 5 mM MgSO_4_, 20 mM imidazole, 10% glycerol, 3 mM β-mercaptoethanol). Proteins were purified by affinity chromatography using the NiNTA agarose (Machery-Nagel), dialyzed against lysis buffer and digested with Ulp1 protease at 4°C overnight. The last purification step was size exclusion chromatography at the HiLoad 16/600 Superdex 200 gel filtration column (GE Healthcare) in SEC buffer (10 mM Tris pH 7.4, 150 mM NaCl, 5% glycerol, 1 mM TCEP). Purified proteins were concentrated to 7 mg/ml and stored in −80°C until needed.

### Preparation of hexamer

*In vitro* transcription was performed in a 50 μL mixture containing: 80 ng/μL of template DNA (6A), 1 mM NTPs (only those necessary for the RNA production), 1.6 mM Np_*n*_Ns, 5% DMSO, 0.12% triton X-100, 12 mM DTT, 4.8 mM MgCl_2_ and 1x reaction buffer for T7 RNAP and 62.5 units of T7 RNAP (New England BioLabs, NEB). The mixture was incubated for 2 h at 37°C. After incubation the samples were injected, without any further purification, in the HPLC and only the hexamer RNA was collected. The purified RNA was dried up on a Speedvac system for three times to remove the excess of Triethylammonium acetate (TEAA).

### *In vitro* transcription with T7 RNAP for 35mer

*In vitro* transcription was performed in a 50 or 75 μL mixture containing: 80 ng/μL of template DNA (35A or 35G), 1 mM NTPs, 1.6 mM Np_*n*_Ns (or ATP or GTP for the control experiments), 5% DMSO, 0.12% triton X-100, 12 mM DTT, 4.8 mM MgCl_2_ and 1x reaction buffer for T7 RNAP and 62.5 units of T7 RNAP (New England BioLabs, NEB). The mixture was incubated for 2 h at 37°C.

### DNAse I treatment

After the transcription, the DNA template was digested by DNAse I to obtain pure RNA. Transcription mixture (50 μL), 6 μL of 10× reaction buffer for DNAse I (10 mM Tris-HCl, 2.5 mM MgCl_2_, 0.5 mM CaCl_2_, pH 7.6 at 25 °C, supplied with the enzyme) and 4 units of DNAse I (NEB) were incubated at 37 °C for 60 min. The enzyme was thermally deactivated at 75 °C for 10 min followed by immediate cooling on ice. All samples were purified with RNA Clean and Concentrator™ from ZYMO research for further use.

### nsp10-nsp16 reaction for screening of the substrates

To test the methyltransferase activity, the cap or the capped-RNA samples were divided into two parts. The positive control contained 200 μM of free cap or ∼40 μM of the RNA (*in vitro* transcribed after DNAse I treatment and purified on RNA Clean and Concentrator™), 1 mM of SAM and 1.5 μM of nsp10/16 in the reaction buffer (40 mM Tris-HCl, 1 mM MgCl2, 5 mM DTT, pH 8 at 25 °C). nsp10-nsp16 was replaced by water for the negative control. The mixture was incubated at 30 °C for 2 h. The enzyme was heat deactivated at 75 °C for 10 min followed by immediate cooling on ice. The reaction with free caps was analyzed without further purification by HPLC and capped-RNA was digested before analysis by LC-MS.

### HPLC Data Collection and Analysis

HPLC was performed using a Waters Acquity HPLC e2695 instrument with PDA detector and with a Kinetex ® XB-C18 column (2.6 μm, 2.1 mm x 50 mm). The mobile phase A was 100 mM TEAA pH 7, and the mobile phase B 100% acetonitrile. The flow rate was kept at 1 mL/min and the mobile phase composition gradient was as follows: linear decrease from 0% to 12% B (6.5% for dimer analysis) over 20 min; linear decrease to 100% B over 7 min; maintaining 100% B for 3 min; returning linearly to 0% B over 10 min. Waters Fraction Collector III was used for collection of the hexamer RNA.

### RNA digestion for LC–MS

The capped-RNA after nsp10-nsp16 reaction was digested using 3 U of Nuclease P1 (Merck) in 50 mM ammonium acetate buffer (pH 4.5) at 37 °C for 1 h. The digested RNA was purified using Amicon-Millipore filters 10 kDa (Merck) to get rid of Nuclease P1. The flow through was dried on a Speedvac system and dissolved in 10 μL of a mixture of acetonitrile (10%) and ammonium acetate (10 mM, pH 9).

### LC–MS data collection and analysis

LC–MS was performed using a Waters Acquity UPLC SYNAPT G2 instrument with an Acquity UPLC BEH Amide column (1.7 μm, 2.1 mm × 150 mm, Waters). The mobile phase A consisted of 10 mM ammonium acetate, pH 9, and the mobile phase B of 100% acetonitrile. The flow rate was kept at 0.25 mL/min and the mobile phase composition gradient was as follows: 80% B for 2 min; linear decrease to 50% B over 4 min; linear decrease to 5% B over 1 min; maintaining 5% B for 2 min; returning linearly to 80% B over 2 min. For the analysis, electrospray ionization (ESI) was used with a capillary voltage of 1.80 kV, a sampling cone voltage of 20.0 V, and an extraction cone voltage of 4.0 V. The source temperature was 120 °C and the desolvation temperature 550 °C, the cone gas flow rate was 50 L/h and the desolvation gas flow rate 250 L/h. The detector was operated in negative ion mode. 8 μL of the dissolved material was injected and analyzed.

### Calculation of methylation efficiency

MassLynx software was used for data analysis and quantification of the relative abundance of all caps. The Area Under the Curve (AUC) for all cap in the positive and negative samples were calculated and normalized for the area of GMP of each negative. The decreasing of the AUC of the starting material (unmethylated cap) in the nsp10-nsp16 treated sample was compared with the AUC of the starting material (unmethylated cap) in the untreated sample and expressed as percentage.

### nsp10-nsp16 reaction for testing of inhibitor

For each reaction ∼10 μM m^7^Gp_3_A-RNA (*in vitro* transcribed, DNAse I treated and purified on RNA Clean and Concentrator™), 100 μM of SAM, 500 nM of nsp10-nsp16 and 50 nM – 5 μM of Sinefungine were added in the reaction buffer (40 mM Tris-HCl, 1 mM MgCl2, 5 mM DTT, pH 8 at 25 °C). The mixtures were incubated at 30 °C for 2 h. The enzyme was heat deactivated at 75 °C for 10 min followed by immediate cooling on ice. The m^7^Gp_3_A-RNA was digested by Nuclease P1 and analyzed by LC-MS.

### LC–MS condition for screening of the nsp10-nsp16 inhibitor

The LC-MS conditions were optimized for the highest signal/noise ratio of m^7^Gp_3_Am RNA cap. LC–MS was performed using a Waters Acquity UPLC SYNAPT G2 instrument with an Acquity UPLC BEH Amide column (1.7 μm, 2.1 mm × 150 mm, Waters). The mobile phase A consisted of 10 mM ammonium acetate, pH 9, and the mobile phase B of 100% acetonitrile. The flow rate was kept at 0.25 mL/min and the mobile phase composition gradient was as follows: 80% B for 2 min; linear decrease to 50% B over 4 min; linear decrease to 5% B over 1 min; maintaining 5% B for 2 min; returning linearly to 80% B over 2 min. For the analysis, electrospray ionization (ESI) was used with a capillary voltage of 2.7 kV, a sampling cone voltage of 30.0 V, and an extraction cone voltage of 3.0 V. The source temperature was 120 °C and the desolvation temperature 500 °C, the cone gas flow rate was 70 L/h and the desolvation gas flow rate 600 L/h. The detector was operated in positive ion mode. 8 μL of the dissolved material was injected and analyzed.

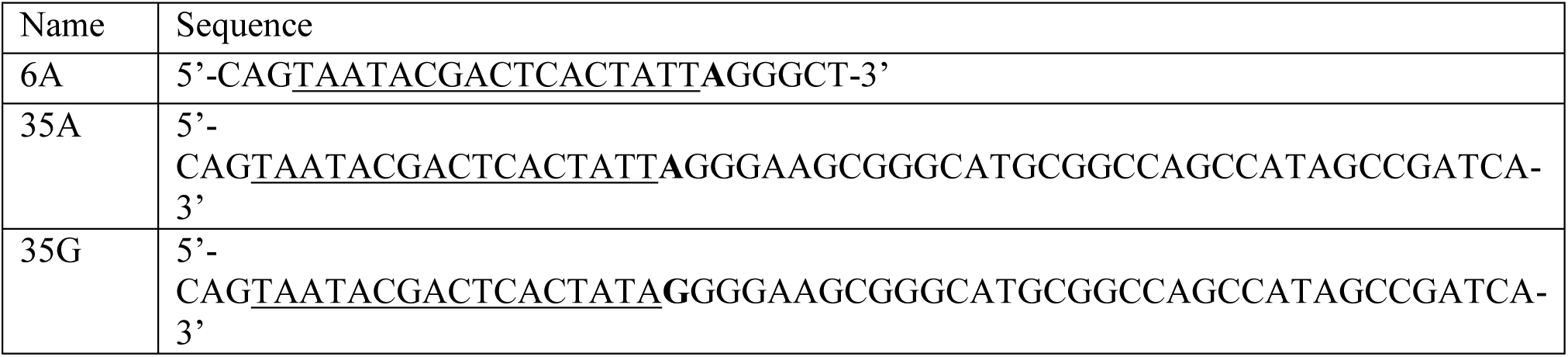

## Author Contributions

RB, PK and MRB performed all experiments; JK, EB, and HC designed and supervised the project; EB and HC wrote the manuscript.

## Competing Interests

The authors declare no competing interests.

## Acknowledgement

The work was supported from European Regional Development Fund; OP RDE; Project: “Chemical biology for drugging undruggable targets (ChemBioDrug)” (No. CZ.02.1.01/0.0/0.0/16_019/0000729), the Academy of Sciences of the Czech Republic (RVO: 61388963) is also acknowledged. We are grateful to Dr. A. Michael Downey (Max Planck Institute of Colloids and Interfaces) for critical reading of the manuscript.

## Supplementary information

### Synthesis and spectroscopic characterization of m^7^GpppA

m^7^GpppA (2160 mOD, 0.101 mmol, 72%) was synthesized by coupling between adenosine 5’-diphospahte triethylammonium salt (ADP) and *N*^7^-methylguanosine 5’-phosphorimidazolide sodium salt (m^7^GMP-Im), which were both prepared as described earlier (Baranowski, J. Org. Chem. 2015, 80, 3982−3997).

ADP (2100 mOD, 0.140 mmol) was mixed with DMSO (0.8 mL) and ZnCl_2_ (228 mg, 1.68 mmol) and left for 10 min under vigorous stirring at room temperature. Then, m^7^GMP-Im (3985 mOD, 0.350 mmol) was added and the reaction progress was monitored by RP HPLC until total conversion of ADP to m^7^GpppA was observed. The reaction was quenched by addition of a solution of Na_2_EDTA (8−10 mmol) and NaHCO_3_ (∼35 mmol) in deionized water (10 ml). The product was purified by DEAE Sephadex chromatography using a linear gradient of triethylammonium bicarbonate buffer (0.9 M) in water. Fractions containing the desired products (as verified by UV, HPLC, and MS analysis) were mixed together and evaporated under reduced pressure with repeated additions of 96% and, then, 99.8% ethanol (to decompose TEAB and remove residual water, respectively). The product was additionally purified by semi-preparative RP HPLC on a VisionHT C18 HighLoad column (Dr. Maisch, 250 mm x 20 mm, 10 μm, flow rate 5 mL/min) using a linear gradient of acetonitrile in 0.05 M ammonium acetate buffer (pH 5.9). The final product was lyophilized three times from water and analyzed by NMR and electrospray MS (ESI-).

^1^H NMR (500 MHz, D_2_O): δ 8.40 (s, 1H), 8.14 (s, 1H), 6.00 (d, *J* = 6.0 Hz, 1H), 5.86 (d, *J* = 3.4 Hz, 1H), 4.65 (t, *J* = 6.0 Hz, 1H), 4.51 – 4.47 (m, 2H), 4.42 – 4.24 (m, 8H), 3.99 (s, 3H); ^31^P NMR (202 MHz, D_2_O) δ −10.35 – −10.89 (m, 2P), −22.20 (t, *J* = 19.4 Hz, 1P); ^31^P NMR {^1^H} (202 MHz, D_2_O) δ −10.63 (d, *J* = 19.3 Hz, 1P), δ −10.67 (d, *J* = 19.3 Hz, 1P), −22.20 (t, *J* = 19.3 Hz, 1P); MS (ESI-)

**Figure S 1:**
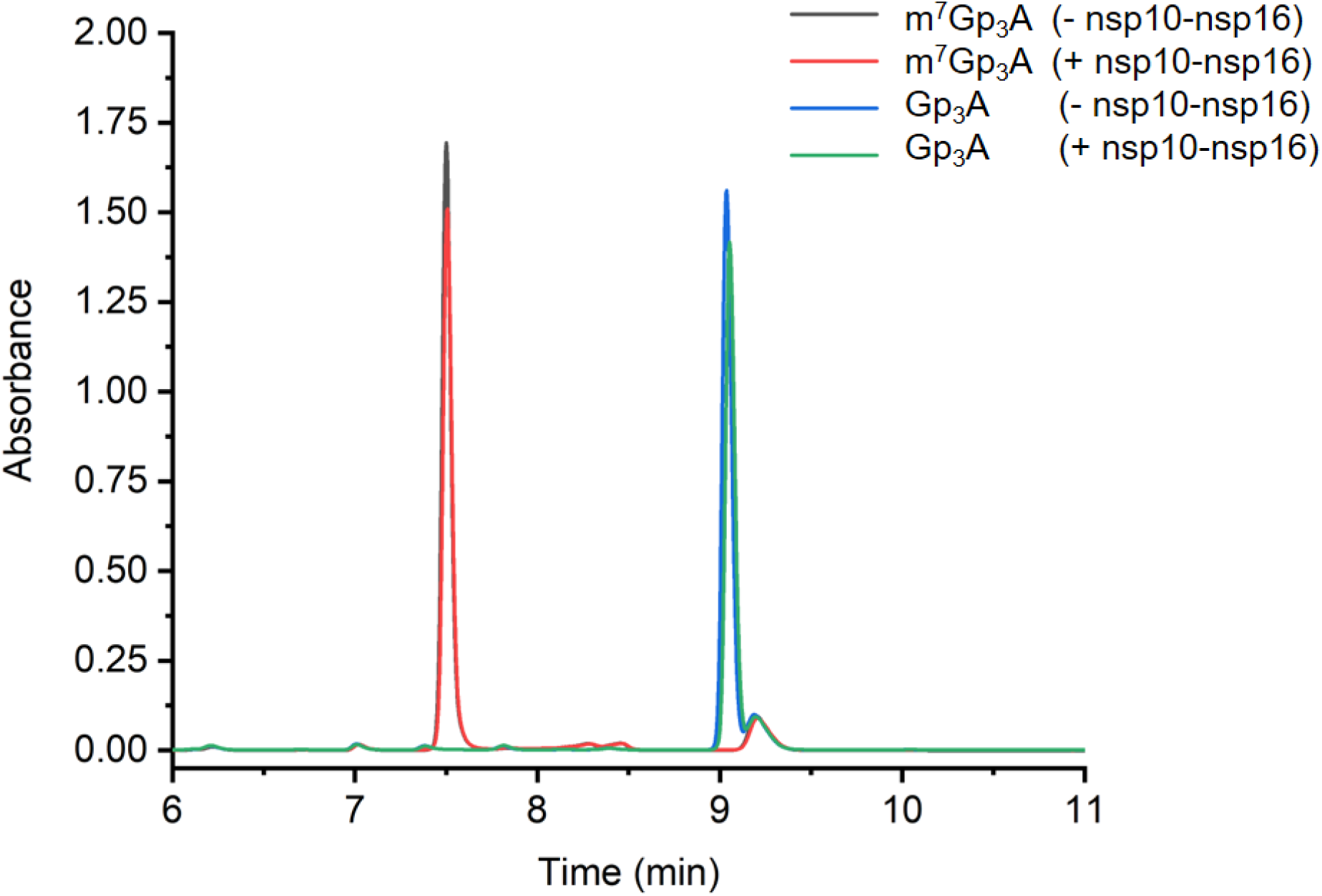
The HPLC chromatogram of free RNA caps before and after the treatement with nsp10-nsp16.

**Figure S 2:**
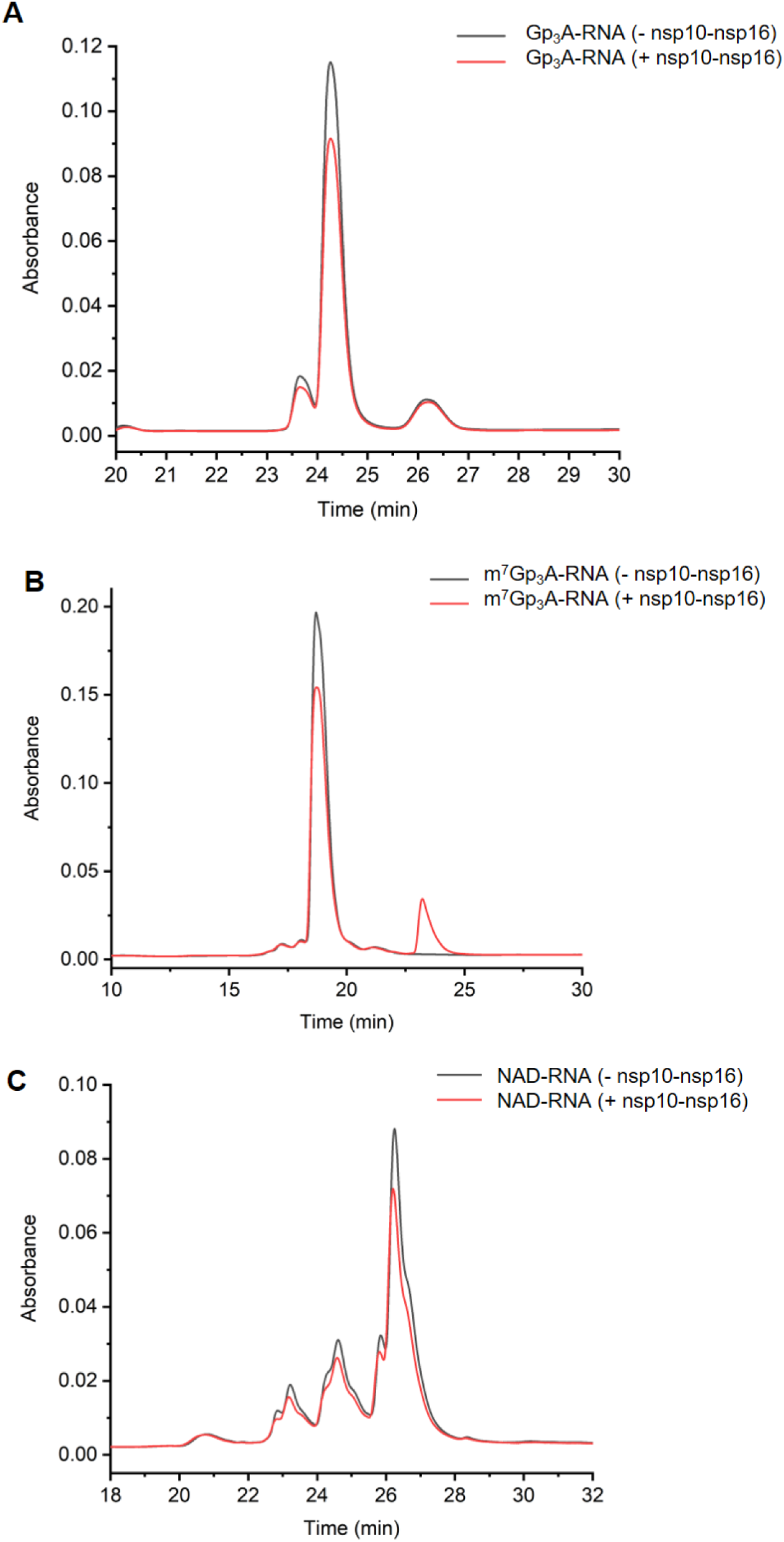
HPLC chromatograms of hexamer RNA capped with Gp_3_A (A), m^7^Gp_3_A (B) and NAD before and after the treatement with nsp10-nsp16.

**Figure S 3:**
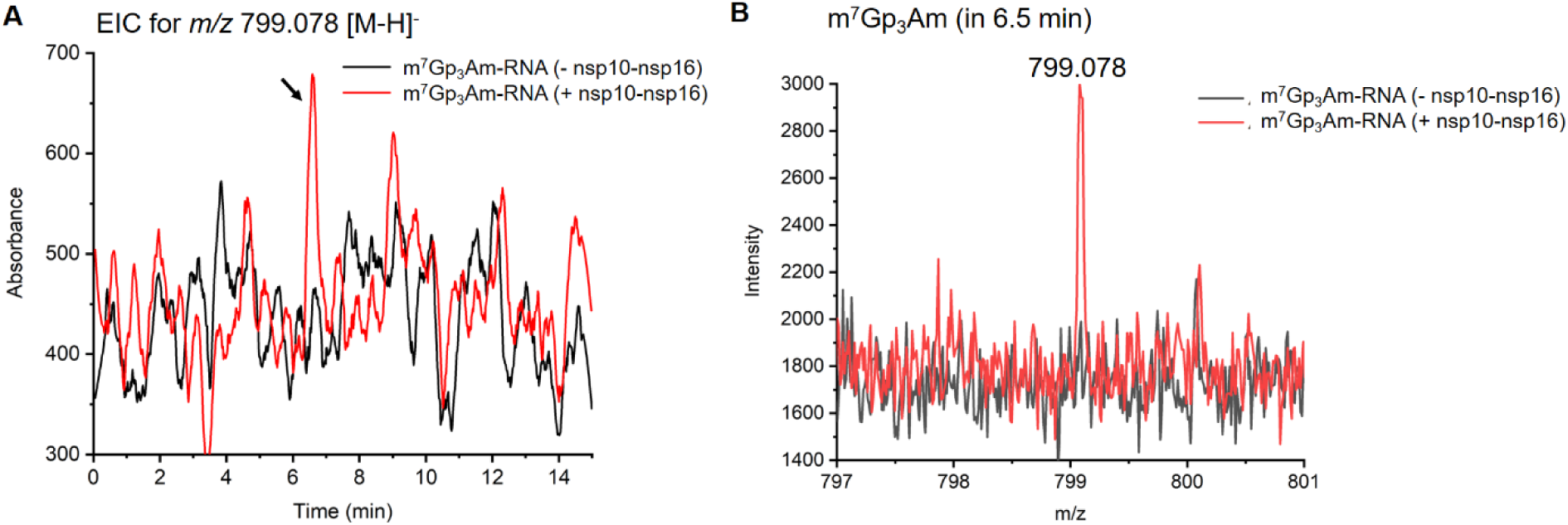
A) Extracted Ion Chromatogram (EIC) of m/z 799.078 from m^7^Gp_3_A-RNA before and after the nsp10-nsp16 treatement. B) MS spectrum of the m/z 799.078 corresponding to m^7^Gp_3_Am before and after the nsp10-bsp16 treatment.

**Figure S 4:**
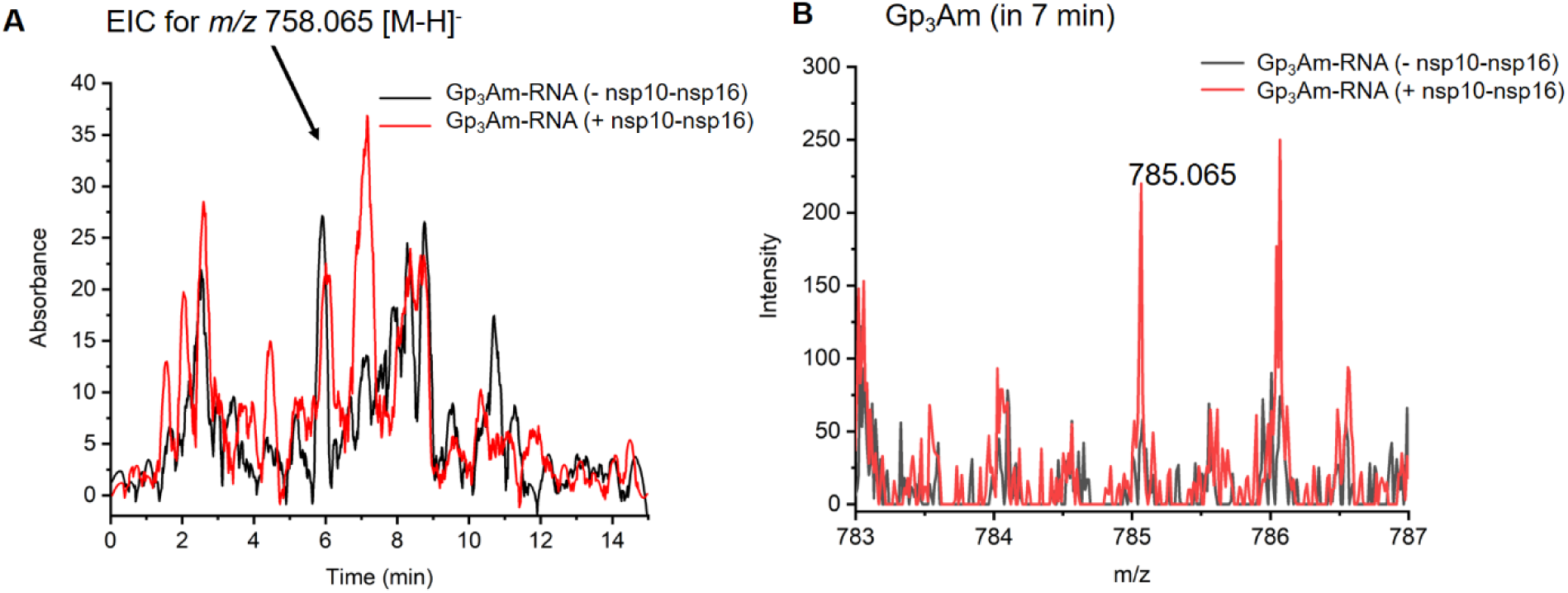
A) Extracted Ion Chromatogram (EIC) of m/z 785.065 from Gp_3_A-RNA before and after the nsp10-nsp16 treatement. B) MS spectrum of the m/z 785.065 corresponding to Gp_3_Am before and after the nsp10-bsp16 treatment.

**Figure S 5:**
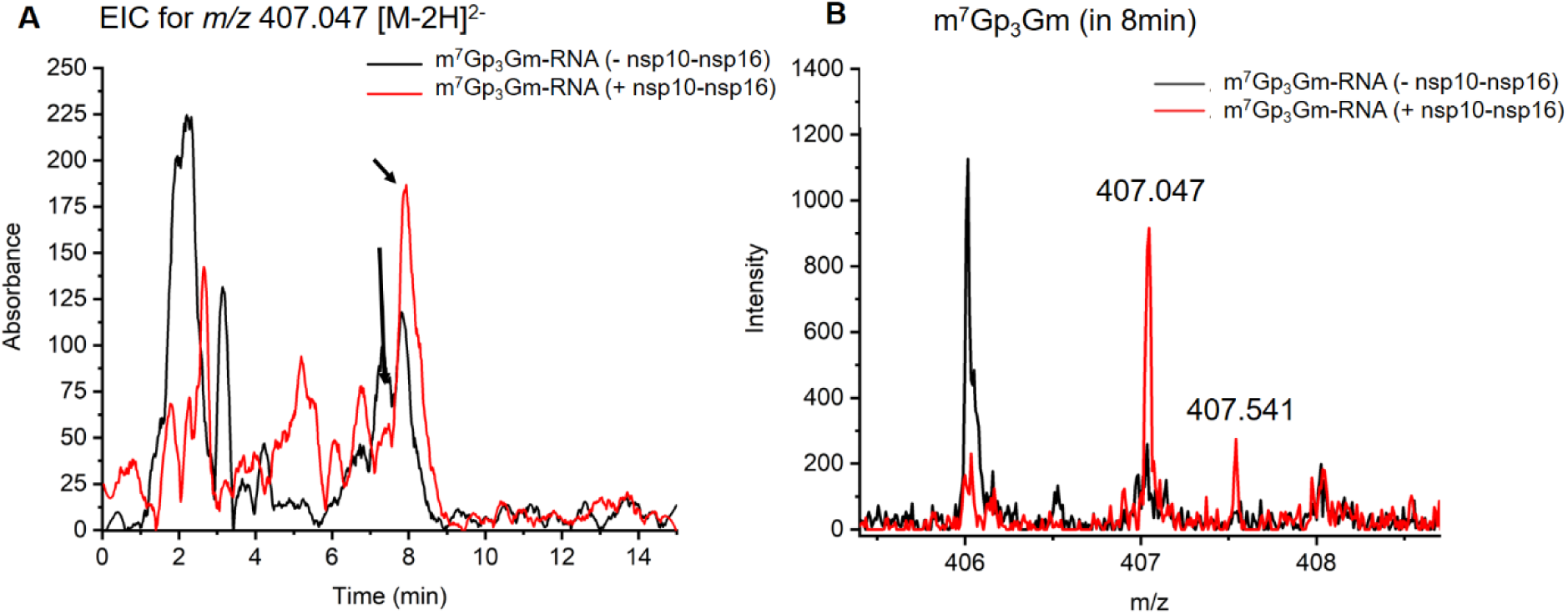
A) Extracted Ion Chromatogram (EIC) of m/z 407.047 from m^7^Gp_3_G-RNA before and after the nsp10-nsp16 treatement. B) MS spectrum of the m/z 407.047 corresponding to m^7^Gp_3_Gm before and after the nsp10-bsp16 treatment.

**Figure S 6:**
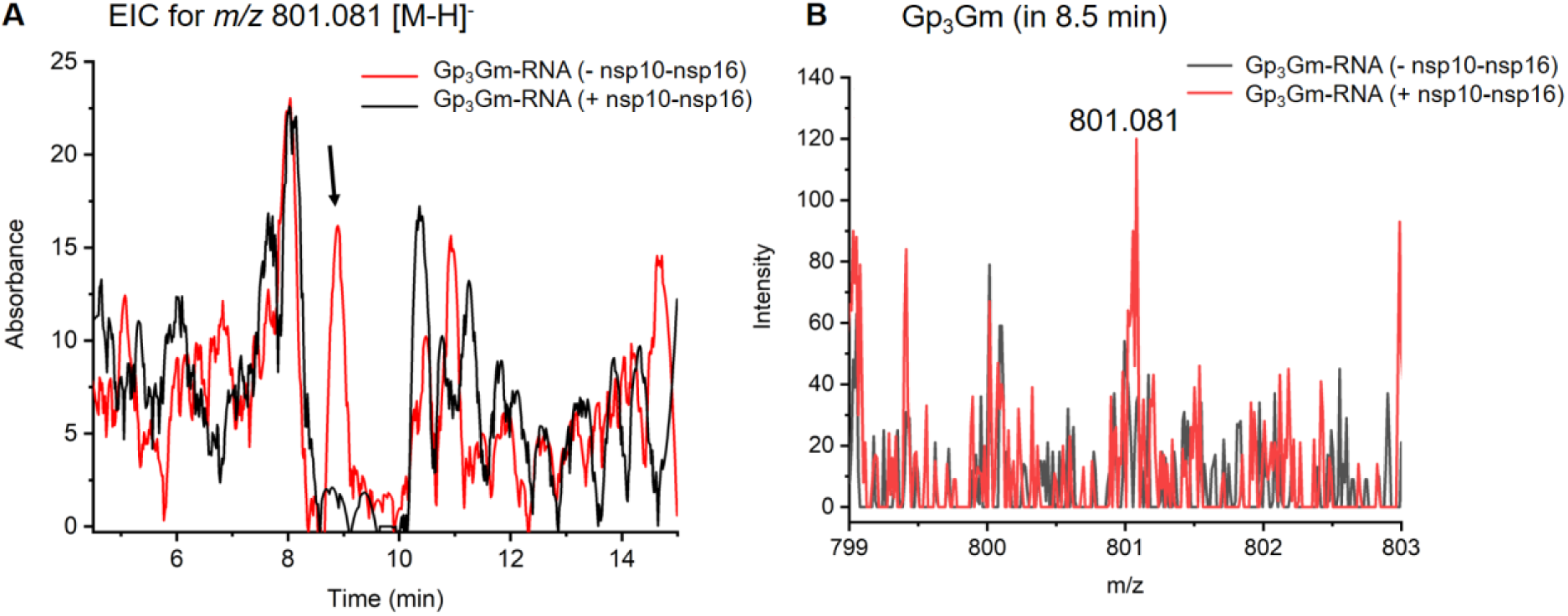
A) Extracted Ion Chromatogram (EIC) of m/z 801.081 from Gp_3_G-RNA before and after the nsp10-nsp16 treatement. B) MS spectrum of the m/z 801.081 corresponding to Gp_3_Gm before and after the nsp10-bsp16 treatment.

**Figure S 7:**
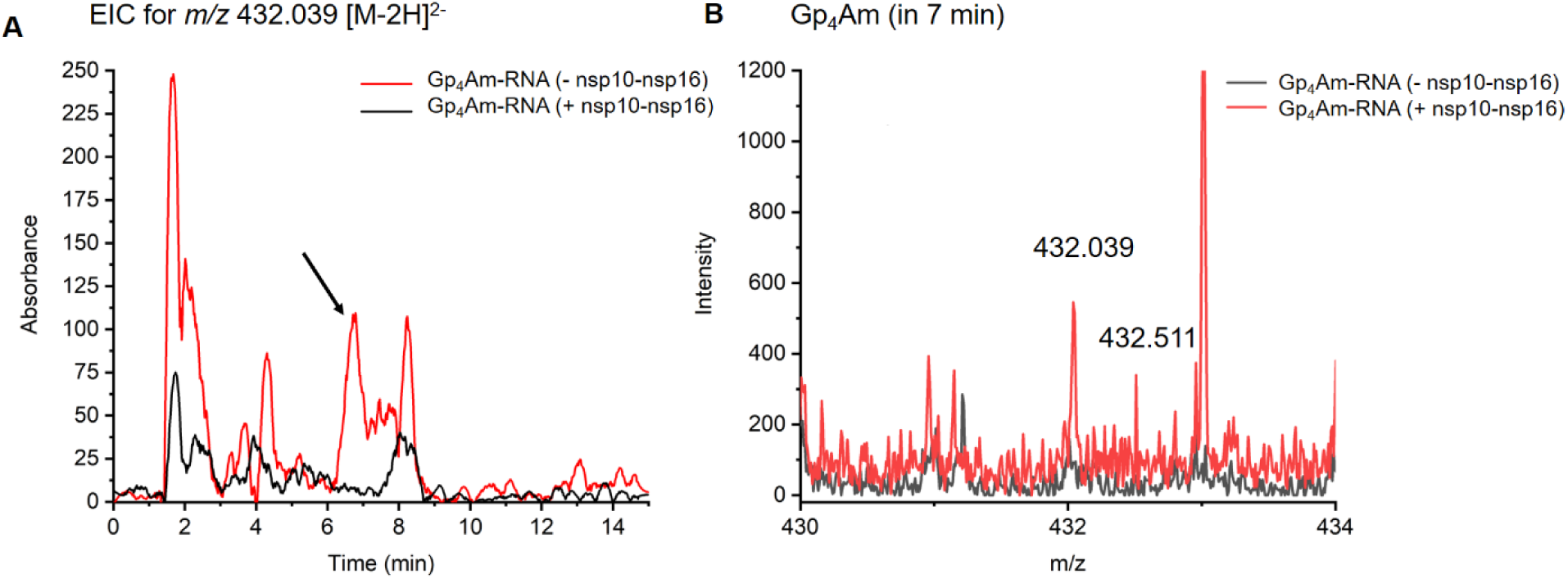
A) Extracted Ion Chromatogram (EIC) of m/z 432.039 from Gp_4_A-RNA before and after the nsp10-nsp16 treatement. B) MS spectrum of the m/z 432.039 corresponding to Gp_4_Am before and after the nsp10-bsp16 treatment.

### NMR spectra

**Spectrum 1.**
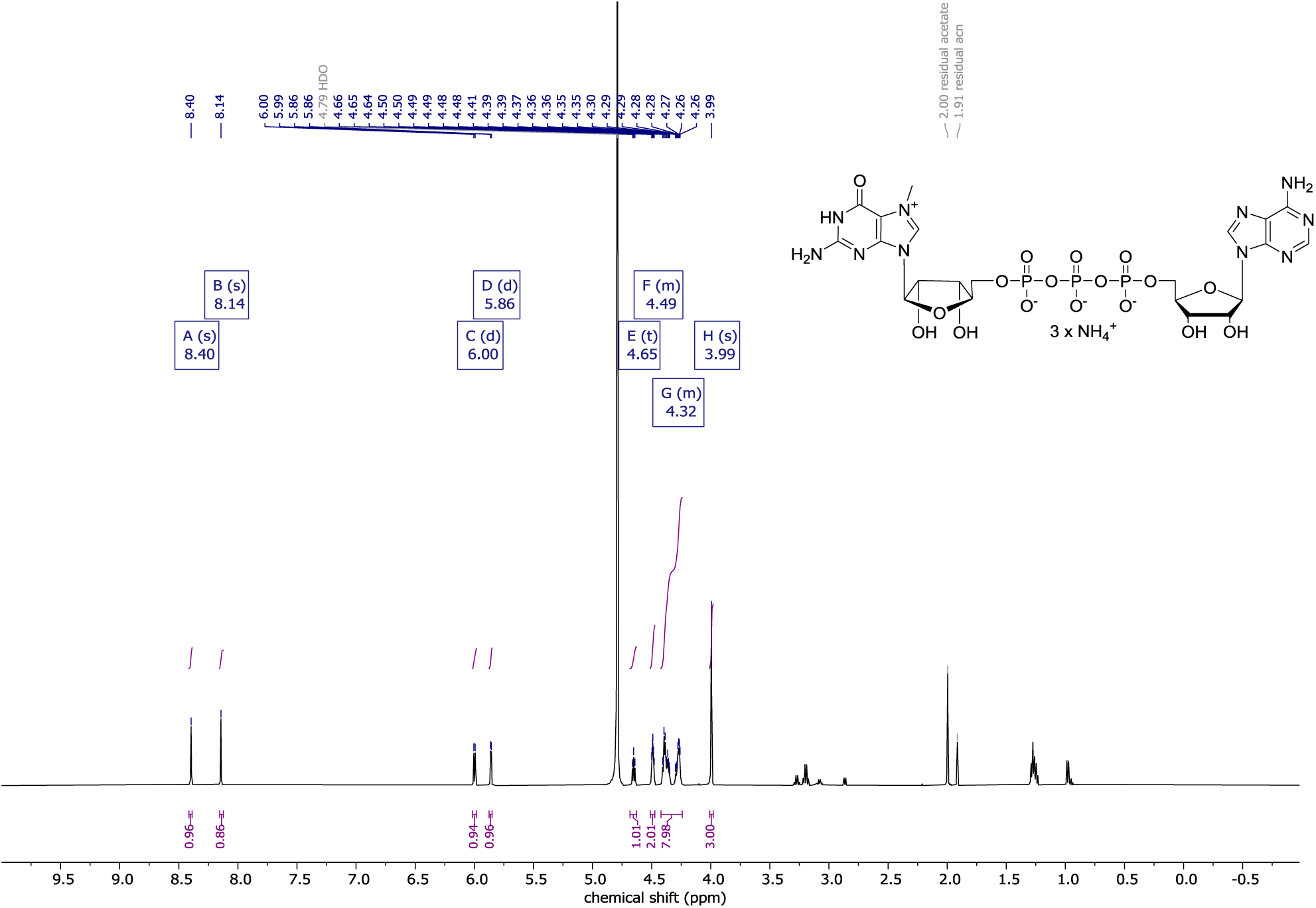
^1^H NMR of m^7^GpppA.

**Spectrum 2.**
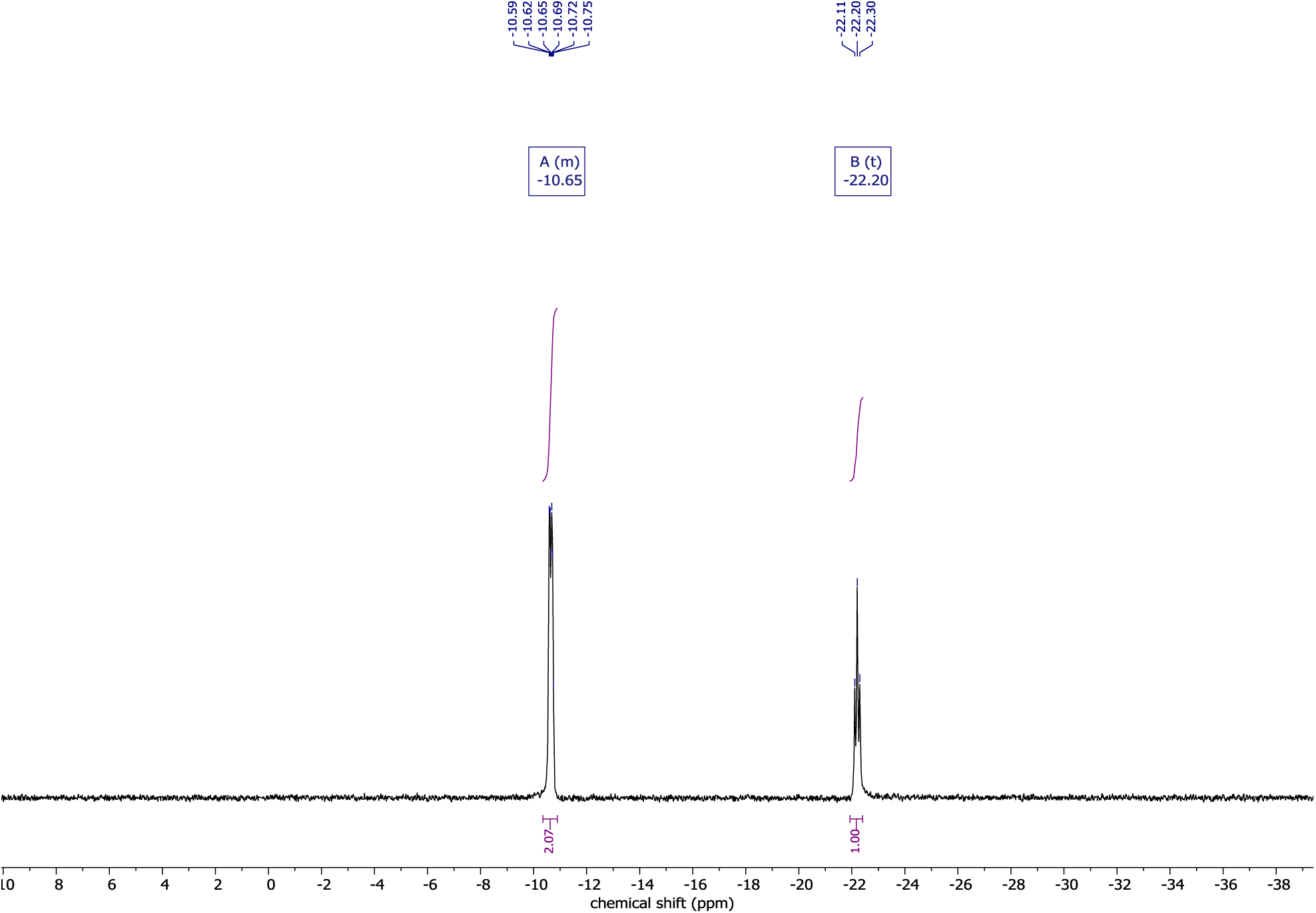
^31^P NMR of m^7^GpppA.

**Spectrum 3.**
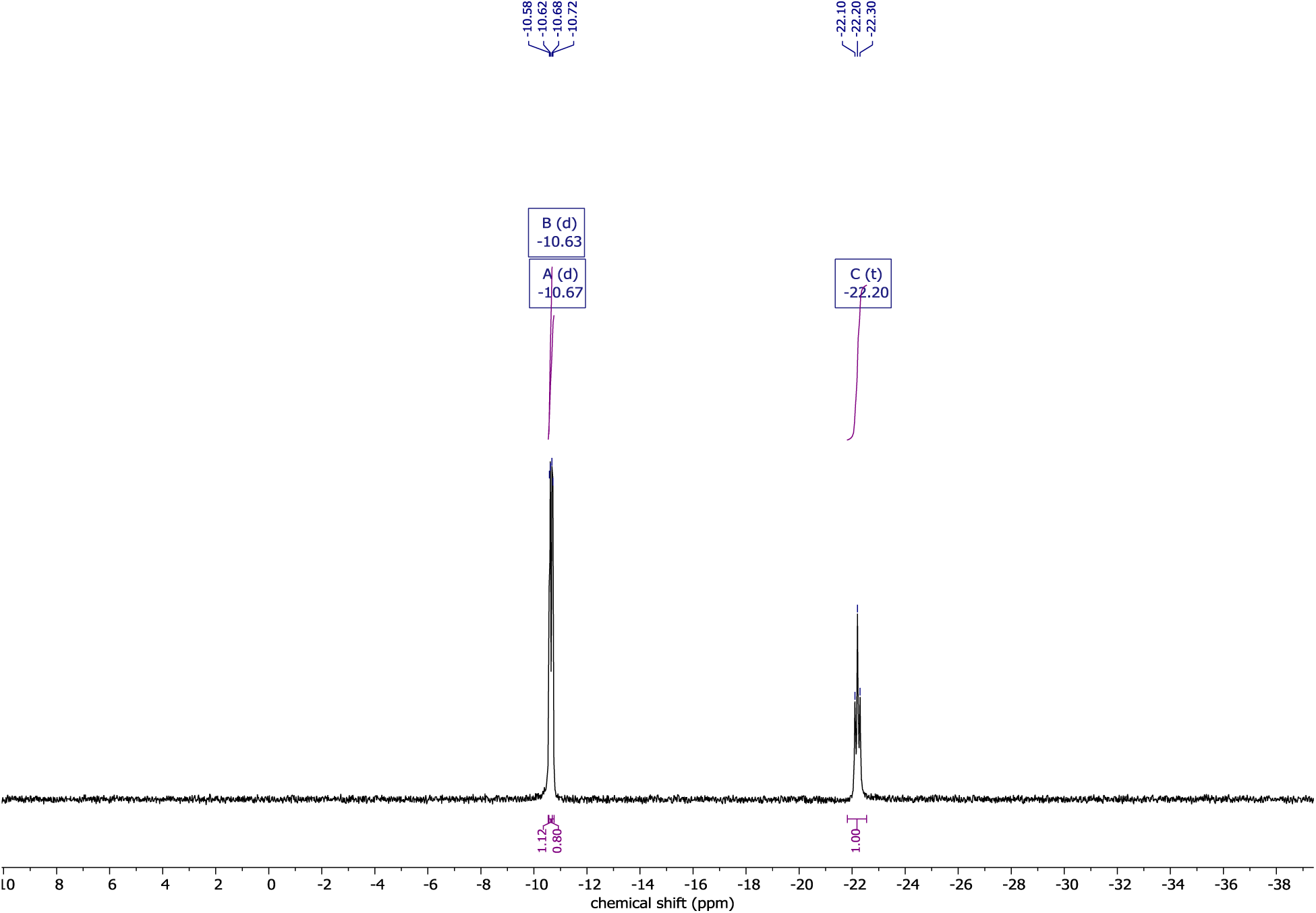
^31^P {^1^H} NMR of m^7^GpppA.

## Notes

### Competing Interest Statement

The authors have declared no competing interest.

